# FastMulRFS: Fast and accurate species tree estimation under generic gene duplication and loss models

**DOI:** 10.1101/835553

**Authors:** Erin K. Molloy, Tandy Warnow

## Abstract

**Motivation:** Species tree estimation is a basic part of biological research but can be challenging because of gene duplication and loss (GDL), which results in genes that can appear more than once in a given genome. All common approaches in phylogenomic studies either reduce available data or are error-prone, and thus, scalable methods that do not discard data and have high accuracy on large heterogeneous datasets are needed.

**Results:** We present FastMulRFS, a polynomial-time method for estimating species trees without knowledge of orthology. We prove that FastMulRFS is statistically consistent under a generic model of GDL when adversarial GDL does not occur. Our extensive simulation study shows that FastMulRFS matches the accuracy of MulRF (which tries to solve the same optimization problem) and has better accuracy than prior methods, including ASTRAL-multi (the only method to date that has been proven statistically consistent under GDL), while being much faster than both methods.

**Availability:** FastMulRFS is available on Github (https://github.com/ekmolloy/fastmulrfs).

## 1 Introduction

Species trees are important models that can be used to address many biological questions, for example how is biodiversity created/maintained and how do species adapt to their environments (Cracraft *et al*., 2002). There is also a vast literature regarding *gene tree reconciliation*, where gene trees are compared to an established species tree in order to understand how genes evolved (for some of the recent literature on this question, see Jacox *et al*., 2016; Lai *et al*., 2017; Muhammad *et al*., 2018; Kundu and Bansal, 2018; Delabre *et al*., 2018; El-Mabrouk and Noutahi, 2019; Dondi *et al*., 2019; Hasić and Tannier, 2019). However, in most cases, species trees are not known in advance and instead must be estimated.

Most species tree estimation methods are designed for orthologous genes, which are genes related through speciation events only and not through duplication events (Fitch, 2000; Moreira and Philippe, 2000). Because orthology prediction is still difficult to do correctly (The Quest for Orthologs Consortium *et al*., 2014; Lafond *et al*., 2018; Altenhoff *et al*., 2019) and mistakes in orthology prediction can result in incorrect species trees, multi-copy genes are often excluded from species tree estimation (e.g., Wickett *et al*., 2014; Leebens-Mack *et al*., 2019). Methods that can estimate species trees from gene families are of increasing interest, as this would enable phylogenetic signal to be extracted from multi-copy genes while avoiding the challenges of orthology prediction.

Several methods have been proposed to infer species trees from multi-copy genes. PHYLDOG (Boussau *et al*., 2013), perhaps the most well-known method explicitly based on a parametric model of gene duplication and loss (GDL), uses likelihood to co-estimate the species tree and gene family trees (which may contain multiple copies from some species). This is very computationally intensive, so PHYLDOG is limited to very small datasets with 10 or so species. Recently, De Oliveira Martins *et al*. (2016) proposed the Bayesian supertree method, *guenomu*, which requires the posterior distribution to be estimated for each gene family tree, for example using MrBayes (Ronquist and Huelsenbeck, 2003). Thus, *guenomu* is also not fast enough to use on genome-scale datasets with 100 or more species.

Non-parametric methods are more commonly used alternatives. For example, gene tree parsimony (GTP) methods take a set of (estimated) gene family trees as input, and then seek a species tree that implies the minimum number of evolutionary events, such as gene duplications and gene losses. Examples of GTP methods include DupTree (Wehe *et al*., 2008), iGTP (Chaudhary *et al*., 2010), and DynaDup (Bayzid and Warnow, 2018). Since GTP is NP-hard, most of these methods operate by using hill-climbing. DynaDup, in contrast, uses dynamic programming to find an optimal solution within a constrained search space; this type of approach, to the best of our knowledge, was first proposed in Hallett and Lagergren (2000) and has since been utilized for other problems, including the maximum quartet support supertree problem (Bryant and Steel, 2001; Mirarab *et al*., 2014) and the Robinson-Foulds supertree problem (Vachaspati and Warnow, 2016). Although GTP methods can be computationally intensive, they are more scalable than other approaches (e.g., PHYLDOG), and several phylogenomic studies have used GTP methods (Sanderson and McMahon, 2007; Burleigh *et al*., 2010).

Other fast approaches include supertree methods that have been adapted to work with gene family trees, referred to as *mul-trees*, as they can have multiple copies from each species. The most well known supertree method for mul-trees is perhaps MulRF (Chaudhary *et al*., 2014b), which attempts to find a solution to the NP-hard Robinson-Foulds Supertree problem for mul-trees (RFS-multree). Although MulRF does not explicitly account for GDL, it has been shown to produce more accurate species trees than DupTree and iGTP on datasets simulated under challenging model conditions with GDL, incomplete lineage sorting (ILS), horizontal gene transfer, and gene tree estimation error (GTEE) (Chaudhary *et al*., 2014a).

In a very recent advance, Legried *et al*. (2020) proved that ASTRAL-multi (Rabiee *et al*., 2019), an extension of ASTRAL (Mirarab *et al*., 2014) to address multi-allele inputs, is statistically consistent under the standard stochastic model of GDL proposed by Arvestad *et al*. (2009) in which all the genes evolve independently and identically distributed (*i.i.d*.) within a species tree, with duplication and loss rates fixed across the edges of the species tree. In fact, ASTRAL-multi is the only method that has been proven statistically consistent under any GDL model. Yet, a comparison reported by Legried *et al*. (2020) between ASTRAL-multi and three earlier species tree estimation methods, including DupTree, STAG (Emms and Kelly, 2018), and MulRF, showed that ASTRAL-multi had good but not exceptional accuracy; specifically, when the duplication and loss rates were both high, ASTRAL-multi was more accurate than DupTree (except when GTEE was low) and STAG (which often failed to complete), but was less accurate than MulRF.

The high accuracy of MulRF in comparison to ASTRAL-multi encouraged us to explore the optimization problem that MulRF attempts to solve (RFS-multree), and led to the following advances.

- We prove (Theorem 5) that the true species tree is an optimal solution to the NP-hard RFS-multree problem, provided there is no adversarial GDL (which occurs when the pattern of duplication and loss events produces bipartitions that are incompatible with the species tree). This model is less restrictive than the standard GDL model in that it does not assume genes evolve *i.i.d*. (similar to the No Common Mechanism model of Tuffley and Steel, 1997), but is more restrictive in that it prohibits adversarial GDL. However, we conjecture (Conjecture 7) that adversarial GDL will occur with sufficiently low probability so that an exact solution to the RFS-multree problem will be statistically consistent for reasonable duplication and loss probabilities.
- We present FastMulRFS, a polynomial-time algorithm that uses dynamic programming to solve the RFS-multree problem exactly within a constrained search space (computed from the input gene family trees), and prove (Theorem 6) that FastMulRFS is statistically consistent under a generic GDL model when no adversarial GDL occurs.
- We prove (Theorem 2) that when solving the RFS-multree problem, any input set of mul-trees can be replaced by a set of smaller trees (with each species labeling at most one leaf), thus reducing memory and running time for methods that attempt to solve the RFS-multree problem.
- We evaluate FastMulRFS in comparison to ASTRAL-multi, DupTree, and MulRF on 1200 diff erent datasets with 100 species and up to 500 genes, generated under 120 model conditions with varying levels of GDL, ILS, and GTEE. We find that FastMulRFS is generally more accurate than DupTree and ASTRAL-multi, and ties for most accurate with MulRF. We also find that FastMulRFS is much faster than MulRF and ASTRAL-multi, and ties for fastest with DupTree. The improvement in performance over ASTRAL-multi is the most important result, as ASTRAL-multi is the only other method to date that has been proven statistically consistent under a stochastic GDL model.

In summary, FastMulRFS is a new and very fast method for species tree estimation that does not require reliable orthology detection and outperforms the leading alternative methods (even under conditions for which FastMulRFS is not yet established to be statistically consistent).

## 2 The RFS-multree problem and FastMulRFS

We define the Robinson-Foulds Supertree problem for mul-trees (RFS-multree) and present FastMulRFS, an algorithm that solves this problem exactly within a constrained search space. Later, we prove that FastMulRFS is statistically consistent under a generic model of GDL when no adversarial GDL occurs. We begin with terminology and definitions.

### 2.1 Terminology

A *phylogenetic tree T* is defined by the triplet (*t, ϕ, S*), where *t* is its unrooted tree topology, *S* is the label set, and *ϕ* : *L*(*t*) → *S* is the assignment of labels to the leaves of *t*. If each label is assigned to at most one leaf, then we say that *T* is *singly-labeled*, whereas if any label is assigned to two or more leaves, then we say that *T* is *multi-labeled* (equivalently, *T* is a *mul-tree*). The edges that are incident with leaves are referred to as *terminal* (or *trivial*) edges, and the remaining edges are referred to as *internal* (or *non-trivial*) edges.

Deleting an edge *e* but not its endpoints from *T* produces two subtrees *t*_*A*_ and *t*_*B*_ that define two label sets: *A* = {*ϕ*(*l*) : *l* ∈ *L*(*t*_*A*_)} and *B* = {*ϕ*(*l*) : *l* ∈ *L*(*t*_*B*_)}. If no label appears on both sides of *e*, then *A* and *B* are disjoint sets, and the edge *e* induces a bipartition *π*_*e*_ on the label set of *T* (i.e., the edge *e* splits the leaf labels into two disjoint sets). However, if some label appears on both sides of *e* then *A* and *B* are not disjoint, and so by definition, the edge *e* does *not* induce a bipartition. We let *C*(*T*) denote the set of bipartitions induced by edges in tree *T*, noting that *not* all edges of *T* will necessarily contribute bipartitions to *C*(*T*), unlike the case of singly-labeled trees.

A key concept in FastMulRFS is *compatibility*, originally described by Estabrook *et al*. (1975), which we now define (see also Warnow, 2017). Let *T*^*^ be the true (fully resolved) species tree on *S*, and let *π* = *A* | *B* be a bipartition on *S*_0_ ⊆ *S*. Then *π* is compatible with *T*^*\**^ if and only if there is a bipartition *π*^′^ = *A*^′^ | *B*^′^ *C*(*T*^*\**^) so that *A* ⊆ *A*^′^ and *B* ⊆ *B*^′^. Equivalently, bipartition *π* on label set *S*_0_ ⊆ *S* is compatible with *T*^*\**^ if there exists *π*^′^ *∈ C*(*T*^*\**^) such that *π*^′^ is identical to *π* when restricted to label set *S*_0_. Similarly, a tree *T* on label set *S*_0_ is compatible with the species tree *T*^*\**^ if every bipartition in *T* is compatible with *T*^*\**^.

### 2.2 Robinson-Foulds Supertree problem for mul-trees

The RF distance (Robinson and Foulds, 1981) between two singly-labeled trees on the same label set has a simple definition as the bipartition distance (i.e., number of bipartitions in one but not in both trees). Now suppose *T* and *T* ^′^ are singly-labeled trees on label sets *S* and *R* ⊆ *S*, respectively. Then the *RF distance* between *T* and *T* ^′^ can be computed as

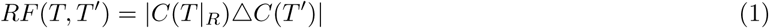

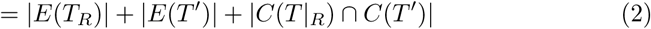

where *T* |_*R*_ denotes *T* restricted to leaves with labels in set *R* (after suppressing internal nodes with degree 2). When one or both trees is a mul-tree, then the RF distance has an alternative definition (which is equal to the standard definition when both trees are singly-labeled and on the same label set): the edit distance under contraction-and-refinement operations, where a contraction is collapsing a single edge, and a refinement is inserting a single edge to decrease the degree of a polytomy (i.e., node of degree four or more). When both trees are mul-trees, computing the RF distance is NP-complete (Chaudhary *et al*., 2013). However, Chaudhary *et al*. (2013) proved that the RF distance between a mul-tree and a singly-labeled tree can be computed in polynomial time as follows: (1) extend *T* with respect to *M*, denoted *Ext*(*T, M*) (Fig. 1), (2) relabel the leaves of *M* and *Ext*(*T, M*) in a *mutually consistent fashion* so that both trees are singly-labeled, and (3) compute the RF distance using Equation 1 between the relabeled versions of *Ext*(*T, M*) and *M*, denoted *Ext*(*T, M*)^′^ and *M* ^′^, respectively; see Appendix for additional details.

**Figure 1:**
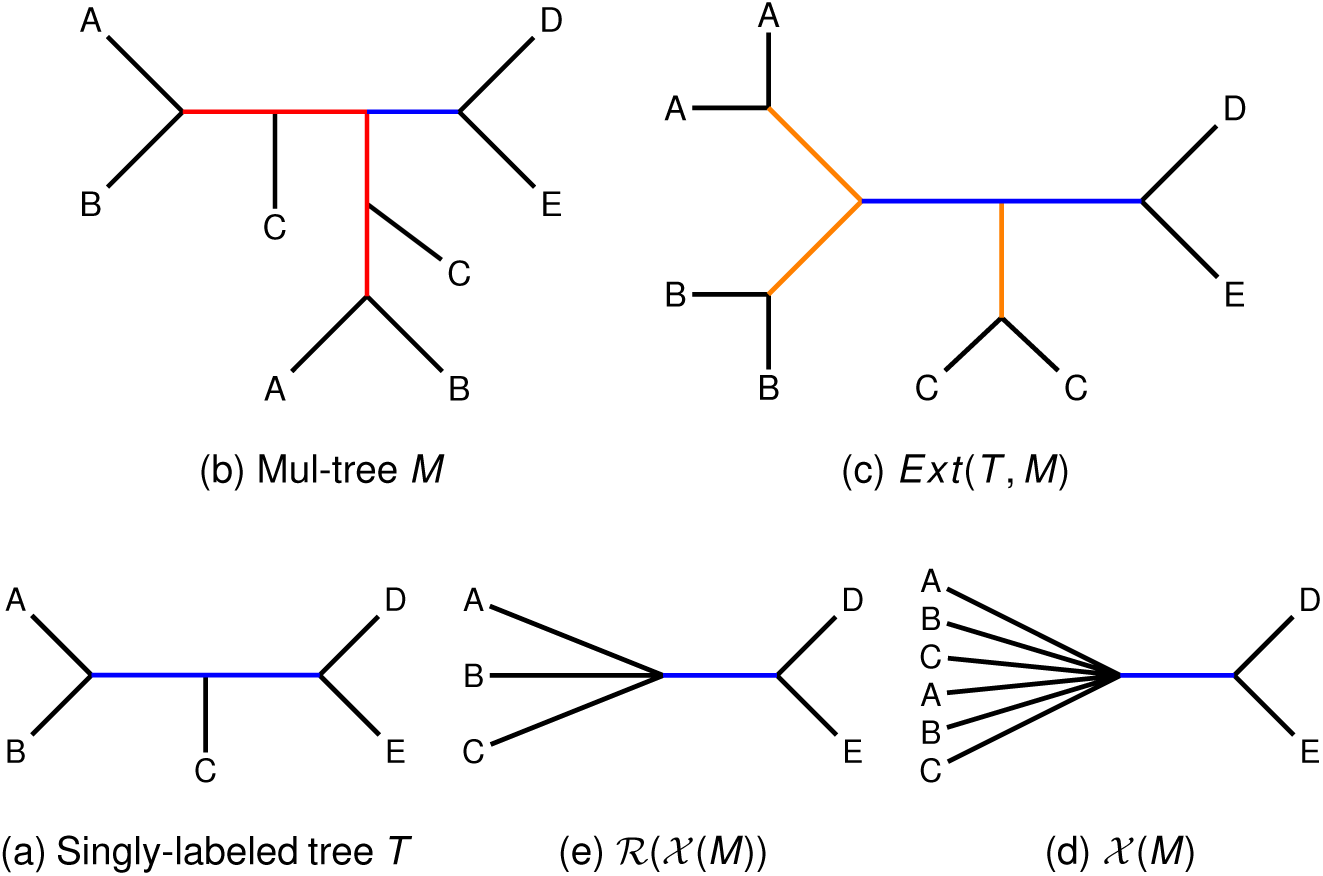
Reduction of the RFS-multree problem to the Robinson-Foulds Supertree (RFS) problem. To compute the RF distance between a singly-labeled tree *T* (subfigure a; bottom-left) and a mul-tree *M* (subfigure b; top-left), we replace *M* by a smaller singly-labeled tree ℛ (𝒳 (*M*)) (subfigure e; bottom-center). We then compute the RF distance between *T* and ℛ (𝒳 (*M*)) using Equation 1. Here we explain why this works. Suppose that *T* (subfigure a) is a candidate singly-labeled, binary supertree for a set *𝒫* of mul-trees and that *M* (subfigure b) is one of the mul-trees in *𝒫*. To compute the RF distance between *T* and *M*, we extend *T* with respect to *M*, producing *Ext*(*T, M*) (subfigure c). Note that *Ext*(*T, M*) has the same non-trivial edges (shown in blue) and the same trivial edges (shown in orange) as *T*, and for every leaf label (species), it has the same number of leaves with that label as mul-tree *M*. The trivial edges in *Ext*(*T, M*) exist in *any possible* singly-labeled, binary tree on *S*; thus, these edges do not impact the solution to the RFS-multree problem. Similarly, mul-tree *M* has edges (shown in red) that will be incompatible with an extended version of *any possible* singly-labeled, binary tree on *S*; thus, these edges do not impact the solution to the RFS-multree problem. An edge is incompatible with every possible singly-labeled supertree if and only if it fails to induce a bipartition (i.e., deleting an edge *e* splits the leaf labels into two non-disjoint sets). Thus, we collapse all internal edges in *M* that fail to induce a bipartition, producing *𝒳* (*M*) (subfigure d). Furthermore, because all leaves with the same label are now on the same side of *every* bipartition in *𝒳* (*M*), we can delete all but one leaf with each label, producing ℛ (*𝒳* (*M*)) (subfigure e). The resulting tree is a non-binary, singly-labeled tree on *S*, so we can compute the RF distance between *T* and ℛ (*𝒳* (*M*)) using Equation 1 when searching for the solution to the RFS-multree problem. These observations are formalized in Lemma 13 (Appendix), and it follows that an RFS-multree supertree for 𝒫 is an RF supertree for 𝒫_*X*_ = {ℛ (*𝒳* (*M*)) : *M*, as summarized in Theorem 2.

Chaudhary *et al*. (2013) then proposed the *Robinson-Foulds Supertree problem for mul-trees* (RFS-multree). The input is a set 𝒫 of mul-trees with leaves labeled by elements of the set *S*, and the output is a binary (i.e., fully resolved) tree *T* bijectively labeled by *S* that minimizes

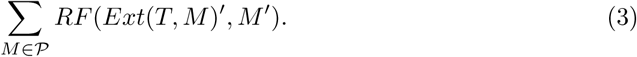

Any tree that minimizes this score is called an **RFS-multree supertree** for 𝒫. Finally, when 𝒫 is a profile of singly-labeled trees, then the RFS-multree problem is the well known Robinson-Foulds Supertree (RFS) problem (Bansal *et al*., 2010; Vachaspati and Warnow, 2016).

### 2.3 Reducing from mul-trees to singly-labeled trees

We simplify the RFS-multree problem by providing an alternative proof that the RF distance between between a singly-labeled tree *T* and a mul-tree *M* and can be computed in polynomial time (Lemma 13, Appendix). We summarize the intuition behind this lemma in Figure 1, which leads easily to Theorem 2.

#### Definition 1

*Given a mul-tree M* ∈ *𝒫, we collapse internal edges with some species labeling leaves on both sides of the edge, denoting the result 𝒳* (*M*). *We then delete all but one leaf with each species label, denoting the result M*_*X*_ = *ℛ* (*𝒳* (*M*)). *We define 𝒫*_*X*_ := {*M*_*X*_ : *M* ∈ *𝒫*}.

#### Theorem 2

*Let T be a singly-labeled, binary tree on label set S, and let 𝒫 be a set of mul-trees. Then, T is an RFS-multree supertree for 𝒫 if and only if T is a RF supertree for 𝒫*_*X*_. *Equivalently, T is an RFS-multree supertree for* 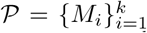 *(with mul-tree M*_*i*_ *on label set S*_*i*_ ⊆ *S) if and only if T is a binary tree that maximizes* 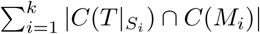.

### 2.4 FastMulRFS

A consequence of Theorem 2 is that any heuristic for the Robinson-Foulds Supertree (RFS) problem can be used for the RFS-multree problem simply by computing *𝒫* _*X*_ (i.e., by transforming the input mul-trees into singly-labeled trees) and then running the heuristic on 𝒫 _*X*_. In this study, we explore the impact of using FastRFS (Vachaspati and Warnow, 2016), an eff ective heuristic for the RFS problem, and we refer to this two-phase approach as **FastMulRFS**.

The input to FastRFS is a profile 𝒯 of singly-labeled trees, each on a (possibly proper) subset of *S* and a set Σ of allowed bipartitions on *S*; FastRFS provably returns a (binary) supertree *T* that minimizes the total RF distance to the trees in 𝒯 subject to *C*(*T*) Σ. FastRFS uses dynamic programming to solve the constrained optimization problem in *O*(*nk* | Σ| ^2^) time, where *n* = | *S*| and *k* =|*𝒯* |. As we will show, Σ can be defined from the input mul-trees so that FastMulRFS runs in polynomial time and is statistically consistent under a generic GDL model when no adversarial GDL occurs.

We now describe FastMulRFS, which takes a profile 𝒫 of mul-trees, each on a (possibly proper) subset of the species set *S*.

1. **Step 1:** We construct *𝒫* _*X*_ from *𝒫* by collapsing all internal edges that have species labeling leaves on both sides of the edge and then deleting all but one of the multiple copies of any species. Thus, *𝒫* _*X*_ is a set of potentially unresolved single-copy gene trees. In the Supplementary Materials (Algorithm 1), we show how to compute the set *𝒫* _*X*_ from *𝒫* in *O*(*mnk*) time, where *n* = |*S*|, *k* = |*𝒫* |, and *m* is the largest number of leaves in any mul-tree in *𝒫*.
2. **Step 2:** We run ASTRAL given the set 𝒫_*X*_ of single-copy gene trees to produce the set Σ of allowed bipartitions. The *default* technique for constructing Σ uses every bipartition in every single-copy gene tree on the *complete* label set *S*. In this case, it is easy to see that |Σ| ≤|{*C*(*M*_*X*_) : *M*_*X*_ *∈ 𝒫*_*X*_ } | ≤ (*n* 3)*k*. Additional bipartitions may be included to guarantee that at least one fully resolved tree *T* satisfies *C*(*T*) ⊆ Σ and to improve accuracy (by expanding the space of allowed solutions); however, ASTRAL-III (Zhang *et al*., 2018) enforces |Σ| = *O*(*nk*). While the total running time of ASTRAL-III is *O*(*nk*|Σ|1.726), we run ASTRAL-III to construct Σ and then exit.
3. **Step 3:** We run FastRFS on the pair (*𝒫*_*X*_, Σ).

In summary, FastMulRFS runs in *O*(*mnk* + *nk* |Σ| ^2^) time, where *n* is the number of species, *k* is the number of mul-trees, and *m* is the largest number of leaves in any of the mul-trees. The default technique for constructing the set Σ of allowed bipartitions enforces |Σ| = *O*(*nk*) and, as we will show in the next section, suffices for proofs of statistical consistency under some generic GDL models.

## 3 Species Tree Estimation using FastMulRFS

### Generic GDL models

Our generic GDL models are similar to the No Common Mechanism models described in Tuffley and Steel (1997), in that there is a common rooted binary model species tree, but each gene evolves down the tree with its own duplication and loss parameters. We make natural assumptions that every gene has duplication probability and loss probability strictly less than 1 on every edge, and note these probabilities can depend on the gene and on the edge. Thus, our generic models contain the GDL models of Arvestad *et al*. (2009) as sub-models.

### Adversarial GDL

We define adversarial GDL to be when the gene evolution process produces a gene family tree with a bipartition *π* that is not compatible with the true species tree *T*^*\**^ (see Section 2.1 for the definition of compatibility). Adversarial GDL requires a sequence of events (a duplication followed by a carefully selected set of losses) that coordinate to produce such a bipartition. Figure 2d illustrates a scenario that produces adversarial GDL: the gene duplicates on the edge above *Y* in the species tree (shown in Figure 2a), so that *Y* has two copies of the gene. Then the first copy of the gene is lost on the edge above *B*, whereas the second copy of the gene is lost on the edge above *A* and the edge above *C*. As a result, the gene family tree shown in Figure 2d is singly-labeled, but the gene family tree induces a bipartition (*A, C* | *B, D*) that is incompatible with the species tree; by definition, this is adversarial GDL. Alternatively, suppose the first copy of the gene had been lost on the edge above *A* and on the edge above (*B, C*), then not only is there no adversarial GDL, but also the gene family tree induces a bipartition (*A, D* | *B, C*) that is compatible with the species tree.

**Figure 2:**
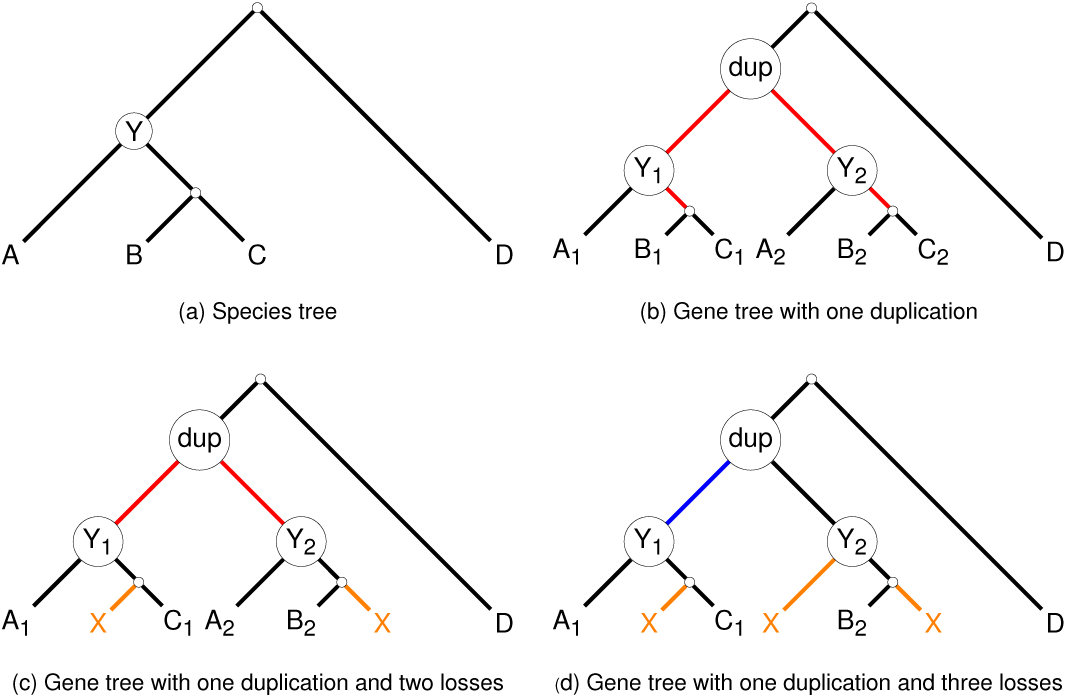
Impact of gene duplications and losses on species tree estimation using RFS-multree methods. Subfigure (a) shows a species tree and subfigures (b) through (d) show three gene family trees that evolved within the species tree. Subfigure (b) shows gene family tree with a duplication event in species *Y* (i.e., the most recent common ancestor of species *A, B*, and *C*). All edges below the duplication (shown in red) fail to induce bipartitions and so will be contracted, and will therefore not impact the solution space for the RFS-multree criterion. Subfigure (c) shows gene tree with a duplication event in species *Y* followed by the first copy of the gene being lost from species *B* and the second copy of the gene being lost from species *C*. Because one of the species that evolved from *Y* retains both copies of the gene, the non-trivial edges below the duplication node fail to induce bipartitions, and so these edges also do not impact the solution space for RFS-multree. Subfigure (d) shows gene family tree with a duplication event in species *Y* followed by the first copy of the gene being lost from species *B* and the second copy of the gene being lost from both species *A* and *C*. None of the species that evolved from *Y* retain both copies of the gene, so all edges below the duplication node induce bipartitions *and hence will not be contracted*; we refer to this situation as “adversarial gene duplication and loss,” because it produces bipartitions in the singly-labeled trees in 𝒫_*X*_ that conflict with the species tree (shown in blue). Such a scenario leads to the possibility that the true species tree may not be an optimal solution to the RFS-multree problem.

Another interesting case to consider is when the gene duplicates on the edge above *Y*, and then the first copy is lost on the edge above *B* and the second copy is lost on the edge above *C*. As a result, the gene family tree shown in Figure 2c does not induce any bipartitions. Now suppose *A, B*, and *C* were clades (rather than leaves), then every edge in the two *A* clades (and the edges connecting the two *A* clades) would fail to induce a bipartition (assuming no other loss events). In contrast, every edge in the *B* clade and the *C* clade would induce a bipartition compatible with the species tree (assuming no other duplication events). In some sense, duplication events hide bipartitions, while losses (following a duplication event) can reveal bipartitions. A carefully selected pattern of losses (after the duplication) can result in adversarial GDL (i.e., a particular bipartition *π* that is not in the species tree), but small changes to that pattern may well produce bipartitions that are in the true species tree or are incompatible with *π*. Thus, overall, while adversarial GDL may occur, it may not have high impact on tree estimation based on the RFS-multree criterion.

In this section, we will discuss model conditions under which adversarial GDL cannot occur: the *duplication-only* case, where all genes evolve with duplication but no loss, and the *loss-only* case, where all genes evolve with loss but no duplication. To prove that a model condition prohibits adversarial GDL, we need to establish that any bipartition that appears in a gene family tree is compatible with the species tree; equivalently, if it appears *in full* in any gene family tree then it must also appear in the species tree, while any incomplete bipartition that appears in any gene family tree can be extended (by adding the missing species) to become a bipartition that is in the species tree. It is trivial to see that if a gene evolves only with losses, then there is no adversarial GDL for that gene (Lemma 3), but the proof for duplication-only evolution is more interesting (Lemma 4).

#### Lemma 3.

*Let 𝒫 be a set of true gene trees that evolved within the rooted species tree T*^*\**^ *under a stochastic loss-only model of gene evolution. Then for π ∈*{*C*(*M*) : *M ∈ 𝒫*}, *π is compatible with T*^*\**^. *Hence, loss-only models have no adversarial GDL*.

#### Lemma 4.

*Let 𝒫 be the set of true gene trees that evolved within the rooted species tree T*^*\**^ *under a stochastic duplication-only model of gene evolution. Then for every mul-tree M ∈ 𝒫, C*(*M*) ⊆ *C*(*T*^*\**^). *Equivalently, for any M ∈ 𝒫, every edge e in M*_*X*_ *(Definition 1) defines a bipartition π*_*e*_ *in C*(*T*^*\**^). *Hence, duplication-only models have no adversarial GDL*.

*Proof*. Let *M* be an unrooted gene family tree, and let *e* be an internal edge in *E*(*M*). We will show that an internal edge *e* is collapsed in producing 𝒳 (*M*) if and only if *e* lies below at least one duplication node in the rooted version of *M*. Hence, the singly-labeled tree *M*_*X*_ = ℛ (𝒳 (*M*)) will only retain the edges in *M* that have no duplication nodes above them in the rooted version of *M*. To see why, consider any edge *e* that has no duplication node above it in the rooted gene family tree: no species appears on both sides of *e* and hence *e* will not be collapsed. Conversely, if internal edge *e* is collapsed, then there must be at least one species on both sides of *e*, and so *e* must be below at least one duplication node in the true rooted gene family tree. Finally, consider a bipartition defined by an edge that is not collapsed, and hence has no duplication nodes above it. This bipartition appears in the true species tree *T*^*\**^, since the only events that cause the gene family tree to diff er from the true species tree are duplications.

We now prove that FastMulRFS is statistically consistent under generic GDL models if no adversarial GDL occurs.

#### Theorem 5

*The true species tree T*^*\**^ *is an RFS-multree supertree for any input 𝒫 for which no adversarial gene duplication and loss occurred*.

*Proof*. The optimization problem seeks a binary tree *T* that minimizes the sum of the RF distances to the input mul-trees; this is equivalent to maximizing the sum of the number of compatible bipartitions in the input mul-trees. If no adversarial GDL occurs, then by definition, every bipartition in the input mul-trees is compatible with the true species tree *T*^*\**^, and so *T*^*\**^ is an optimal solution to the RFS-multree problem.□

#### Theorem 6

*FastMulRFS is statistically consistent under any GDL model for which adversarial GDL is prohibited*.

*Proof*. Let *T*^*\**^ be the true species tree. By Theorem 5, *T*^*\**^ is an optimal solution to the RFS-multree problem for any input 𝒫 for which no adversarial GDL occurred. Since our generic GDL models assume that the probability of no duplication or loss occurring on an edge is always strictly positive for every gene, the true species tree has strictly positive probability of appearing in the set 𝒫 of gene family trees. Therefore, as the number of genes increases, Σ (as constructed by the default setting within FastMulRFS) will converge to *C*(*T*^*\**^) with probability converging to 1, and *T*^*\**^ will be the unique tree that is optimal under the RFS-multree problem for input *𝒫*. FastMulRFS finds an optimal solution to RFS-multree problem subject to the tree *T* it returns satisfying *C*(*T*) *⊆* Σ, by Theorem 2 and by Theorem 3 in Vachaspati and Warnow (2016). Since Σ converges to *C*(*T*^*\**^) as the number of genes increases, the probability that FastMulRFS will return *T*^*\**^ converges to 1. □

We finish this section with a conjecture.

#### Conjecture 7

*FastMulRFS is statistically consistent under a generic model of GDL for probabilities of gene duplication and loss, so that adversarial GDL has sufficiently low probability*.

## 4 Experimental Study

We evaluated FastMulRFS in comparison to ASTRAL-multi, DupTree, and MulRF on biological and simulated datasets, considering species tree topological accuracy and running time. All simulated datasets are available on the Illinois Data Bank (https://doi.org/10.13012/B2IDB-5721322_V1), and the commands necessary to reproduce this study are provided in the Supplementary Materials.

### Biological dataset

We analyzed a fungal dataset with 16 species and 5351 genes from Rasmussen and Kellis (2012), who provided gene family trees estimated from their nucleotide alignments. In a prior study, Butler *et al*. (2009) estimated species trees from this same dataset (specifically the concatenated amino acid alignment of putatively orthologous sequences) using MrBayes (Ronquist and Huelsenbeck, 2003), constrained to enforce the out-grouping of *S. castellii* with respect to *S. cerevisiae* and *C. glabrata*. The other reported trees diff ered with respect to this group (i.e., not all analyses returned this as a clade) and diff ered in the placement of *K. waltii*. According to their study, none of these resolutions are clearly correct.

### Simulation study

We generated a collection of 100-species datasets (each with 1000 model gene trees) under the DLCoal model (Rasmussen and Kellis, 2012), which is a unified model of GDL and ILS. The easiest model condition was based on parameters estimated from the 16-species fungal dataset (Rasmussen and Kellis, 2012; Du *et al*., 2019), and then we increased the GDL rates and ILS levels (by increasing population size) to make more challenging model conditions. We used RAxML (Stamatakis, 2014) to estimate gene trees under the GTR+г model from the simulated alignments, with sequence lengths varied to produce four diff erent levels of GTEE. Finally, we estimated species tree giving methods the first 25, 50, 100, and 500 gene family trees, either true or estimated, as input. This created 120 model conditions (3 GDL rates, 2 levels of ILS, 5 levels of GTEE, and 4 numbers of genes), each with 10 replicates, for a total of 1200 datasets. Importantly, none of the model conditions prohibits adversarial GDL, allowing us to explore method performance when adversarial GDL may occur.

### Evaluation criteria

On the fungal biological dataset, we evaluated accuracy with respect to established evolutionary relationships, and on the simulated datasets, we quantified error using the RF error rate, with respect to the true (model) species tree. We also recorded empirical running time; however, it should be noted that all experiments were performed on the Campus Cluster at the University of Illinois at Urbana-Champaign, which is a heterogeneous system (i.e., compute nodes do not have the same specifications; see here: https://campuscluster.illinois.edu/resources/docs/nodes/).

## 4.1 Results

### Results on biological dataset

We analyzed the fungal dataset using ASTRAL-multi, FastMulRFS, DupTree, and MulRF. All produced trees that are very similar to the MrBayes concatenation tree (Supplementary Figure S1), and the diff erences are minor given (1) the variability in the trees found by Butler *et al*. (2009), (2) the use of a topological constraint in their MrBayes analysis, and (3) the uncertainty about the placement of specific taxa in the tree. For further information on these analyses, see Section 5 in the Supplementary Information from Butler *et al*. (2009).

Given that the topological diff erences are minor, we report the running time diff erences. FastMulRFS and DupTree completed in under a minute each, ASTRAL-multi completed in 18 minutes, and MulRF completed in 40 minutes. Hence, FastMulRFS is much faster than MulRF and ASTRAL-multi. While all four of these methods are relatively fast on 16 taxa, we expect the diff erence between methods to increase on datasets with larger numbers of species and higher rates of gene duplication. The improvement in running time over MulRF and ASTRAL-multi is due in part to the fact that both MulRF and ASTRAL-multi use the original gene family trees, while FastMulRFS uses the reduced singly-labeled trees; hence, as the number of leaves or the duplication rate increase, the advantage in running time for FastMulRFS should also increase.

### Results on the simulated datasets

DupTree had poorer accuracy than the other tested methods (Section 4.1 in Supplementary Materials). Hence, we focus on comparing MulRF, FastMulRFS, and ASTRAL-multi. The fastest method was FastMulRFS, MulRF was the slowest, and ASTRAL-multi was intermediate. All methods improved in accuracy with larger numbers of genes and degraded in accuracy with higher GTEE levels, ILS levels, and/or GDL rates. The relative accuracy between methods was consistent across all model conditions, although the degree of diff erence depended on the model conditions, with bigger diff erences for smaller numbers of genes and higher GTEE levels, ILS levels, and GDL rates. When given 500 gene trees, error levels were low and diff erences between methods were (usually) small, so that the main diff erence was running time. We present results in Figure 3 for MulRF, FastMulRFS, and ASTRAL-multi under the highest GDL rate, the highest level of ILS, and the second highest level of GTEE (about 53%). We note that high GTEE (such as in this setting) is consistent with the generally low bootstrap branch support values reported for several phylogenomic datasets (e.g., about 25% for exon and 45% for intron datasets from Jarvis *et al*., 2014; also see Table 1 in Molloy and Warnow, 2018). See Supplementary Materials for additional results.

**Figure 3:**
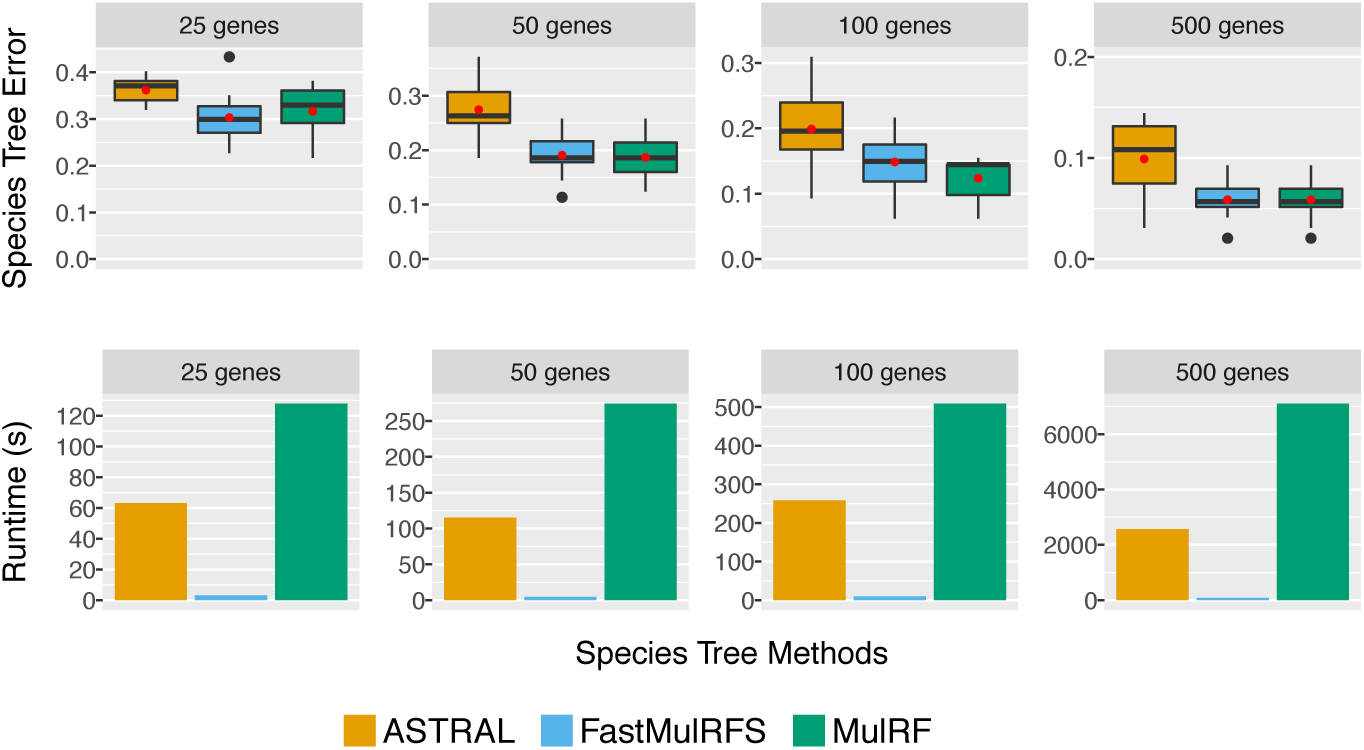
Species tree error rate (i.e., RF error rate) and running time (seconds) are shown for FastMulRFS, MulRF, and ASTRAL-multi under the most challenging model conditions with 100 species. All datasets have the second highest GTEE level (moderate GTEE: 52%), the highest ILS level (low/moderate ILS: 12%), and the highest GDL rate (D/L rate: 5 × 10^7^). Red dots (first row) and bars (second row) are means for 10 replicate datasets.

### FastMulRFS vs. MulRF

Both try to solve the RFS-multree problem but use diff erent approaches; they were essentially tied for accuracy across all tested conditions, but FastMulRFS was dramatically faster (Fig. 3; Supplementary Tables S3–S4). In addition, FastMulRFS nearly always returned trees with better RFS-multree scores than MulRF (Section 4.2 in Supplementary Materials).

### FastMulRFS vs. ASTRAL-multi

Figure 3 shows results for the second highest GTEE level, where FastMulRFS was much more accurate than ASTRAL-multi for all numbers of genes. FastMulRFS was always at least as accurate as ASTRAL-multi (often more accurate) across the other model conditions (Supplementary Table S3), with larger diff erences between methods for the higher GTEE condition and smaller diff erences for the lower GTEE conditions. The running times for ASTRAL-multi and FastMulRFS increased with the number of genes, but FastMulRFS was always much faster (Fig. 3, Supplementary Table S4). For example, on the 500-gene model conditions, FastMulRFS typically completed in 1–2 minutes (and always in under 5 minutes), but ASTRAL-multi used between 10 minutes and 1.2 hours.

## 5 Discussion

To date, only two methods have been proven statistically consistent under any GDL model— ASTRAL-multi and FastMulRFS—but the conditions under which these two methods have been proven statistically consistent are diff erent. ASTRAL-multi is established consistent under a gene evolution model that allows both gene duplication and loss to occur for each gene, but requires that all the genes evolve *i.i.d*.. In contrast, FastMulRFS has been proven consistent under a generic model that does not require the genes to evolve *i.i.d*. (and indeed allows for a very broad no-common-mechanism model); this is a relative strength for the theoretical result for FastMulRFS, as genes do not evolve *i.i.d*. down a species tree, as discussed in Dondi *et al*. (2019). On the other hand, FastMulRFS has only been proven consistent when no adversarial GDL occurs; this is a relative weakness of the theoretical result for FastMulRFS (although see Conjecture 7). Thus, from a theoretical perspective, there are advantages and disadvantages for both methods.

We now consider the empirical performance of the methods evaluated in this study, focusing on the simulated datasets (since diff erences on the biological dataset were minor, except for running time). Under most of the model conditions we examined, FastMulRFS was more accurate and more robust to GTEE than ASTRAL-multi. Furthermore, the only conditions in which the two methods achieved similar accuracy were characterized by low GTEE and large numbers of genes, where both methods achieved very high accuracy. In addition, FastMulRFS was much faster than ASTRAL-multi, with large improvements in speed, especially for large numbers of genes and high GTEE. Thus, FastMulRFS had superior performance compared to ASTRAL-multi, the only previous method to date established statistically consistent under a stochastic GDL model.

A comparison between FastMulRFS and MulRF is also interesting. Both methods attempt to solve the same NP-hard optimization problem, and neither is guaranteed to find an optimal solution. However, FastMulRFS is guaranteed to find an optimal solution within a constrained search space within polynomial time, whereas MulRF uses a local search strategy that is not guaranteed to find optimal solutions and is not guaranteed to complete in polynomial time. Furthermore, the way that FastMulRFS constrains its search space is sufficient to ensure that it is statistically consistent, but this statement is not guaranteed for MulRF. From a theoretical perspective, therefore, FastMulRFS is superior to MulRF. In terms of empirical performance in our study, the two methods were very close in accuracy, but FastMulRFS was dramatically faster. Therefore, overall, FastMulRFS was superior to MulRF.

We note that FastMulRFS matched or improved on the other methods under all conditions we explored, where gene trees evolved under a unified model of ILS and GDL (which did not prohibit adversarial GDL). Hence, our study suggests that FastMulRFS may have good robustness and high accuracy, even under conditions where it has not (yet) been proven statistically consistent. However, future work is clearly needed to evaluate FastMulRFS and other methods under a wider range of model conditions, including explicit conditions where adversarial GDL occurs.

## 6 Summary and Conclusions

FastMulRFS is a new method that can estimate species tree from unrooted gene family trees, without needing to have any information about orthology. FastMulRFS is provably statistically consistent under a GDL model that allows genes to evolve under a no-common-mechanism model (a more general model than the Arvestad *et al*. (2009) *i.i.d*. model assumed in the proof of statistical consistency for ASTRAL-multi), provided that adversarial GDL does not occur. Prior to this study, ASTRAL-multi was the only method proven to be statistically consistent for estimating species trees in the presence of GDL.

FastMulRFS always matched or improved on the accuracy of ASTRAL-multi (often substantially) in our simulation study, which included three GDL, two ILS levels, and five GTEE levels, and it was also faster than ASTRAL-multi. Furthermore, these model conditions do not prohibit adversarial GDL. This improvement in accuracy over ASTRAL-multi is significant, since our proof only establishes statistical consistency under models where no adversarial GDL, ILS, or GTEE is present. Although accuracy is difficult to evaluate on biological datasets, FastMulRFS produced trees that were similar to those produced by other methods and did not violate known relationships.

This study suggests several directions for future work. In particular, we should explore additional simulation conditions to evaluate the impact of higher GDL rates (including conditions that explicitly have adversarial GDL) and larger numbers of genes, where the relative performance of species tree estimation methods might be different. Simulations should also be performed to evaluate other scenarios that produce multi-copy genes, for example whole genome duplication events, which impact species tree estimation for many major clades, including fungi (Butler *et al*., 2009) and plants (Leebens-Mack *et al*., 2019). More complex simulations should also be considered, including ILS, introgression, gene conversion, etc., in order to better understand the conditions in which each method performs well. Furthermore, it would be helpful to characterize biological datasets in understand realistic levels of ILS and GDL (including the frequency of adversarial GDL).

A limitation of this study is that we only examined a few methods, and future studies should also evaluate other methods, including *guenomu* (discussed earlier) and MixTreEM (Ullah *et al*., 2015), to discover the places in the parameter space of model species trees where each method outperforms the others. Furthermore, methods that operate by making predictions of orthology could be used in a three-phase approach: given inputs with sequence alignments and MUL-trees, predict orthology, reduce to datasets with just orthologous genes (and hence singly-labeled gene trees), and then run a preferred species tree estimation method. For example, in a recent preprint, Zhang *et al*. (2019) presented another modification of ASTRAL, A-PRO, and proved it statistically consistent under a GDL model if given correctly rooted and “tagged” gene trees (i.e., each node in each gene tree is correctly identified as either a duplication or a speciation); however, this assumption means that orthology can be inferred without error (an assumption that is not made for ASTRAL-multi). Future studies should evaluate A-PRO as well in estimating a species tree from MUL-trees. Such studies would enable biologists to select methods with the best expected accuracy for their datasets.

An important direction for future work is to evaluate the theoretical properties (such as statistical consistency) of FastMulRFS under parametric GDL models, where adversarial GDL is possible. The statistical consistency of DupTree and other methods (e.g., MixTrEm, *guenomu*, and even modifications to concatenation to enable such analyses on multi-copy gene family datasets) should also be evaluated.

Overall, the recent advances in development of statistically consistent methods for species tree estimation under GDL models is exciting, and the good performance of many of these methods under a range of model conditions suggests that novel combinations and ideas may lead to even better methods that provide improved accuracy and scalability.

## Supporting information

revised supplementary materials

## Acknowledgements

The authors thank the anonymous reviewers as well as Siavash Mirarab and the members of the Warnow Lab for feedback that improved this paper.

### Appendix

We begin with the following two additional definitions from Ganapathy *et al*. (2006); Chaudhary *et al*. (2013).

#### Definition 8 (Full Differentiation)

*We say that M* ^′^ = (*m, ϕ*^′^, *S*^′^) *is a full diff erentiation of mul-tree M* = (*m, ϕ, S*) *if ϕ*^′^ : *L*(*m*) → *S*^′^ *is a bijection. In other words, M* ^′^ *is a singly-labeled version of M*.

#### Definition 9 (Mutually Consistent Full Diff erentiations)

*Let* 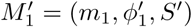 *and* 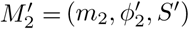 *be full diff erentiations of mul-trees M*_1_ = (*m*_1_, *ϕ*_1_, *S*) *and M*_2_ = (*m*_2_, *ϕ*_2_, *S*), *respectively. For i* = 1, 2, *we define R*_*i*_(*s*) *⊆ S*^′^ *to be the set of labels given to the leaves in* 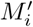 *that are labeled s in M*_*i*_. *We say that* 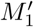 *and* 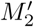 *are mutually consistent full diff erentiations (MCFDs) of M*_1_ *and M*_2_ *if R*_1_(*s*) = *R*_2_(*s*) *8s* ∈ *S*.

Ganapathy *et al*. (2006) showed that if *M*_1_ and *M*_2_ are both mul-trees, then their RF distance can be computed as

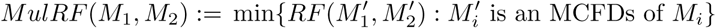

which implies an exponential-time algorithm for computing the RF distance between two mul-trees (Ganapathy *et al*., 2006), Later, Chaudhary *et al*. (2013) showed this problem is NP-complete and introduced a special case, where one of the two mul-trees has the property: every leaf with the same label is grouped together into polytomy that is separated by an edge from the rest of the tree. A mul-tree with this property can be viewed as an extended version of a singly-labeled tree.

#### Definition 10 (Extended Version)

*Let T* = (*t, ϕ*_*T*_, *S*) *be a singly-labeled tree, and let M* = (*m, ϕ*_*M*_, *S*) *be a mul-tree. Let k*_*s*_ *be the number of leaves with label s in M. The extended version of T with respect to M, denoted Ext*(*T, M*), *is created by attaching k*_*s*_ *new leaves to the leaf labeled s in T, assigning label s to each of these new leaves, and repeating this process for all s* ∈ *S*.

Chaudhary *et al*. (2013) showed that the RF distance between a mul-tree *M* and (the extended version of) a singly-labeled tree *T*, both on label set *S*, can computed in polynomial time. Here, we provide an alternative proof that further simplifies this problem. First, we present two transformations that can be applied to a mul-tree *M* = (*m, ϕ, S*) or to its full diff erentiation *M* ^′^ = (*m, ϕ*^′^, *S*^′^) by using the function *f* : *S*^′^ → *S* with property that *f* (*ϕ*^′^(*l*)) = *ϕ*(*l*) for all *l* ∈ *L*(*m*).

#### Definition 11 (Contracted Version)

*The contracted version of M, denoted 𝒳* (*M*), *is created by contracting every edge e that fails to induce a bipartition, because some species label appears on both sides of e. Similarly, the contracted version of M* ^′^, *denoted 𝒳* (*M* ^′^), *is created by contracting every edge e with π*_*e*_ = *A*|*B such that f*(*A*) ∩ *f*(*B*) ≠ ø.

#### Definition 12 (Reduced Version)

*If all leaves with species label s are on the same side of* every *edge in E*(*m*), *then they can be represented by a single leaf labeled s. The reduced version of M or M* ^′^, *denoted ℛ* (*M*) *or ℛ* (*M* ^′^), *respectively, is created as follows. For every s ∈S with the aforementioned property, delete all but one of the leaves in the set* {*l ∈L*(*m*) : *f* (*ϕ*^′^(*l*)) = *ϕ*(*l*) = *s*} *(suppressing internal vertices of degree 2) and relabel the remaining leaf s*.

It is easy to see that ℛ (𝒳 (*M* ^′^)) is a singly-labeled tree that is isomorphic to ℛ (*𝒳* (*M*)), because after applying the function 𝒳 to either *M* ^′^ or *M*, all the leaves with species label *s* will be on the same side of every edge and thus can be replaced by a single leaf with species label *s* by applying the function *R*. This observation holds for all *s* ∈ *S*.

#### Lemma 13

*Let T be a singly-labeled, fully resolved tree on label set S, let M* = (*m, ϕ, S*) *be a mul-tree, and let Ext*(*T, M*)^′^ *and M* ^′^ = (*m, ϕ*^′^, *S*^′^) *be MCFDs of Ext*(*T, M*) *and M, respectively. Then*,

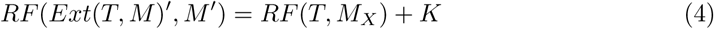

*where M*_*X*_ = ℛ (*𝒳* (*M*)) *and K is a constant that does not depend on the topology of the singly-labeled tree T on S*.

*Proof*. Let *f* : *S*^′^*→S* be a function with property that *f* (*ϕ*^′^(*l*)) = *ϕ*(*l*) for all *l ∈ L*(*m*), and define *X* = {*A*| *B∈C*(*M* ^′^) : *f* (*A*) *f* (*B*) ≠ } ø and *R* = *A B C*(*M* ^′^) : *A >* 1, *B >* 1, and either *f* (*A*) = 1 or *f* (*B*) = 1. Thus, *X* contains bipartitions that *cannot exist* in *C*(*Ext*(*T, M*)) for any singly-labeled tree *T* on *S*, and *R* contains bipartitions that *must exist* in *C*(*Ext*(*T, M*)) for any singly-labeled tree *T* on *S*. Let *E*^′^ denote *Ext*(*T, M*)^′^. Then,

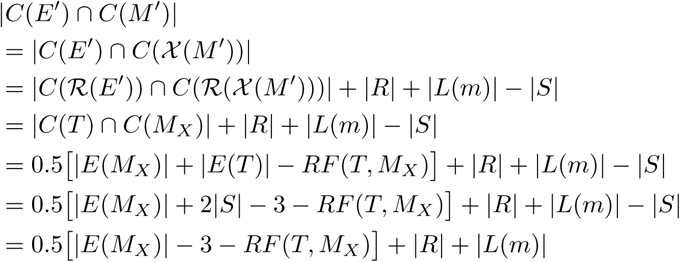

Let *c* be the number of species in *M* that have multiple copies. Then,

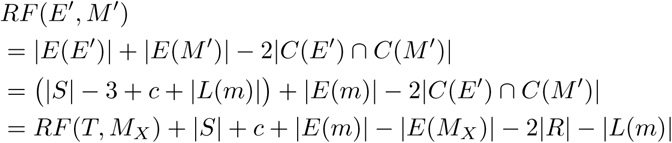

where *S, c, E*(*m*), *E*(*M*_*X*_), *R*, and *L*(*m*) are independent of *T*.

In Lemma 13, we show that *MulRF* (*Ext*(*T, M*),*M*) can be computed in polynomial time, because computing *RF* (*Ext*(*T, M*)^′^,*M* ^′^) does not depend on the MCFDs of *Ext*(*T, M*) and *M*. In addition, we show that the RF distance between *Ext*(*T, M*) and *M* can be computed (up to a constant factor that does not depend on the topology of a singly-labeled tree *T* on *S*) by simply transforming *M* into a (potentially unresolved) singly-labeled tree on *S* and computing its RF distance from *T*. This gives us Theorem 2.

