## Supplementary material for "FastMulRFS: Fast and accurate species tree estimation under generic gene duplication and loss models": revised supplementary materials

May 7, 2020

#### Contents

**List of Tables** 1

**List of Figures** 1

**1 Algorithm for Preprocessing Mul-trees** 2

**2 Results on Biological Dataset** 3

**3 Simulation Study** 3

**References** 11

### 1 Algorithm for Preprocessing Mul-trees

**Algorithm 1:** Reduction on MUL-trees.

```
Input   : MUL-tree T = (t, φ, S) with |L(t)| = m and |S| = n
Output: R(X(T))

Function ReduceMultree(T):
    T ← RootArbitrarily(T)
    below ← [0]_{2m×n}; above ← [0]_{2m×n}; found ← [0]_n

    // Step 1:  Fill vector with 1 if species is below node and 0 otherwise.
    for v ∈ PostOrderNodeTraversal(T) do
        if IsLeaf(v) then
            below[v, φ(v)] ← 1
        else
            l → GetLeftChild(v); r → GetRightChild(v)
            for s ∈ S do
                below[v, s] ← below[l, s] ∨ below[r, s]

    // Step 2:  Fill vector with 1 if species is "above" node and 0 otherwise.
    root ← GetRoot(T)
    l ← GetLeftChild(root); r ← GetRightChild(root)
    for s ∈ S do
        above[l][s] ← below[r][s]
        above[r][s] ← below[l][s]
    for v ∈ PreOrderNodeTraversal(T) do
        if v ≠ root and v ≠ l and v ≠ r and not IsLeaf(v) then
            p ← GetParent(v); x ← GetLeftChild(p)
            if v == x then
                x ← GetRightChild(p)
            for s ∈ S do
                above[v][s] ← above[p][s] ∨ below[x][s]

    // Step 3:  Contract edges that fail to induce bipartitions.
    for v ∈ PostOrderInternalNodeTraversal(T) do
        p ← GetParent(v)
        SetEdgeLength((v, p), 1)
        for s ∈ S do
            if below[v][s] ∧ above[v][s] then
                SetEdgeLength((v, p), 0)
                Break
    ContractEdgesWithZeroLength (T)

    // Step 4:  Remove extra copies of species.
    for l ∈ L(t) do
        s ← φ(l)
        if found[s] then
            RemoveLeaf(l)
        else
            found[s] ← 1

    return Unroot(T)
```

### 2 Results on Biological Dataset

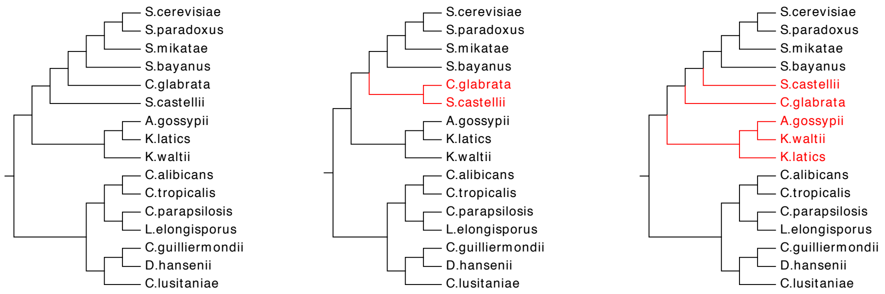

(a) Concatenation analysis (b) ASTRAL-multi (c) FastMulRFS and other methods

Figure S1: Species trees estimated on the the 16-taxon fungi dataset from ]Species trees estimated on 16-taxon fungi dataset with 5,351 estimated gene trees from [1]. The concatenation analysis tree (a) is based on putative orthologs instead of the multi-gene family trees, the ASTRAL-multi tree on the gene family trees is shown in (b), and "FastMulRFS and other methods" (c) refers to the tree found in common by MulRF, DupTree, and FastMulRFS. The topological differences between the trees are fairly small, and the true placement of these taxa is not established. DupTree and FastMulRFS were the fastest (both completed in less than a minute), ASTRAL-multi estimated a species tree in 18 minutes, and MulRF completed in 40 minutes.

### 3 Simulation Study

All datasets and scripts used in this study are available on the Illinois Data Bank (`https://doi.org/10.13012/B2IDB-5721322_V1`).

#### 3.1 SimPhy Simulation

Here we describe the simulation of gene trees from a species tree under a model of GDL. Our protocol is based on the simulation study by [1] that used a biological dataset (16 species of fungi) from [6]. We simulated gene trees with different rates of GDL and ILS based on the species tree estimated by Rasmussen and Kellis [6] (download: `http://compbio.mit.edu/dlcoal/pub/config/fungi.stree`), which had estimated branch lengths (in millions of years). We multiplied each branch length by $10^7$ (i.e., assuming 10 generations per year) to get a tree height of 1,800,000,337.5 generations. We then found (through visualization of 16-taxon species trees) that speciation rates of $1.8 \times 10^{-10}$ and $1.8 \times 10^{-8}$ corresponded to deep and recent speciation, respectively; we used the intermediate value of $1.8 \times 10^{-9}$.
Finally, we ran SimPhy [5] Version 1.0.2 with the command:

```
simphy-1.0.2-mac64 -rs 10 -rl F:$ngen -rg 1 -sb F:$sprt -st $trln \
    -sl F:$ntax -si F:1 -sp F:$psiz -su F:$murt -lb F:$dlrt \
    -ld F:lb -hg LN:1.5,1 -o <output directory> -ot 0 -om 1 -od 1 \
    -op 1 -oc 1 -ol 1 -v 3 -cs 293745 &> <log file>
```

where `$ntax` is the number of taxa (100), `$ngen` is the number of genes (1000), `$trln` is the tree length (1,800,000,337.5 generations), `$sprt` is the speciation rate ($1.8 \times 10^{-9}$ events per generation), `$psiz` is the effective population size (either $1 \times 10^7$ or $5 \times 10^7$), `$murt` is the mutation rate (0.0000000004 mutations per generation per site) and `$dlrt` is the duplication and loss rate (either $1 \times 10^{-10}$, $2 \times 10^{-10}$, or $5 \times 10^{-10}$ events per generation per lineage). Note that the tree-wide effective population size (`-sp`), the tree-wide substitution rate (`-su`), the duplication rate (`-lb`), and the loss rate (`-ld`) are the same parameters used by [1]; these parameters (with GDL rate of $1 \times 10^{-10}$ and effective population size of $1 \times 10^7$) are similar to those estimated from the biological dataset by [6]. We allowed gene trees to deviate from a strict molecular clock by using gene-by-lineage-specific rate heterogeneity modifiers (`-hg`), meaning that a gamma distribution was defined for each gene tree by drawing $\alpha$ from a log-normal distribution with a location of 1.5 and a scale of 1 (same parameters as used in [10]), and then each branch length in a gene tree is multiplied by a value drawn the gamma distribution corresponding to that gene tree.

SimPhy simulates gene trees from a species tree under the unified model of GDL and ILS proposed by Rasmussen and Kellis [6]. This simulation procedure has two steps: first, a collection of locus trees are simulated from the species tree, and second, a gene tree is simulated for each of the locus trees. Gene trees can differ from the species tree due to GDL as well as ILS, whereas a gene tree differs from its locus tree due to ILS *only*. We quantified the level of ILS by computing the normalized Robinson-Foulds (RF) distance [7] between each true locus tree and its respective true gene tree (which are on the same leaf set), averaging this value across all 1000 locus/gene trees. The average locus-to-gene tree discord (AD) across the 10 replicate datasets was 2% and 12% for datasets with an effective population size of $1 \times 10^7$ and 12% for $5 \times 10^7$, respectively. We also quantified the level of GDL by examining the number of leaves and the number of species per gene tree. All gene trees had approximately 100 leaves, as the duplication and loss rates are equal. As the duplication/loss rate increases, the number of species per gene tree decreases, and thus, even though locus/gene trees have the same number of leaves on average, these leaves are labeled by fewer species. For duplication/loss rates of $1 \times 10^{-10}$, $2 \times 10^{-10}$, and $5 \times 10^{-10}$ the average number of species per gene tree was 85, 74, and 53 species; information regarding the number of copies per species is reported in Table S1.

Unlike in the simulation performed by Du *et al.* [1], we did not enable gene conversion and specified 10 generations per year instead of $1.\bar{1}$ generations per year (note that we accidentally used the value estimated for the fly dataset instead of the fungal dataset; thanks to Celine Scornavacca for pointing this out to us!). Fortunately, this parameter choice seems reasonable enough given that Rasmussen and Kellis [6] considered values of 0.6 generations per year up to 10 generations per year for the fungal dataset and selected $1.\bar{1}$ generations per year, as this value resulted in a level of ILS in simulations that was similar to the level of ILS estimated for the fungal dataset.

To characterize the impact of our parameter choice, we simulated datasets specifying $1.\bar{1}$ generations per year (by setting the species tree height `$trln` to be $2 \times 10^8$ generations and the speciation rate `$sprt` to be $2 \times 10^{-8}$ events per generation). This resulted in increased ILS and reduced numbers of duplications/losses, as expected. Specifically, AD was 22% (vs. 2%) and 64% (vs. 12%) for datasets with an effective population size of $1 \times 10^7$ and 12% for $5 \times 10^7$, respectively. The average number of species per gene tree was 98 (vs. 85), 96 (vs. 74), and 90 (vs. 53) for duplication/loss rates of $1 \times 10^{-10}$, $2 \times 10^{-10}$, and $5 \times 10^{-10}$, respectively (note that information regarding the number of copies per species is reported in Table S1). The simulation specifying $1.\bar{1}$ generations per year does not seem realistic given the number of copies of each species in the fungal dataset (Table S2). Consequently, it may be preferable to specify 10 generations per year to increase the species tree height and then increase the population size to evaluate higher levels of ILS. A more problematic issue is whether the DLCoal model is a good fit for the fungal dataset, and future research should explore this issue further and evaluate methods on datasets simulated under different models that result in multi-copy genes.

#### 3.2 INDELible Simulation

Here we describe the simulation of multiple sequence alignments for each model gene tree produced by SimPhy under the GTR+Γ model of evolution. Again, our protocol is also based on the fungal dataset from [6] (`http://compbio.mit.edu/dlcoal/pub/data/real-fungi.tar.gz`), which included a multiple sequence alignment estimated using MUSCLE [2] and a maximum likelihood tree estimated using PhyML [4] for each of the 5,351 genes. We estimated GTR+Γ model parameters for each of the PhyML gene trees by running RAxML Version 8.2.12 [8] with the following command:

```
raxmlHPC-SSE3 -m GTRGAMMA -f e -t <PhyML gene tree file> -s <MUSCLE alignment file> \
            -n <output name>
```

We then fit distributions to the GTR+Γ model parameters estimated from alignments with at least 500 distinct alignment patterns and at most 25% gaps. Finally, we drew GTR+Γ model parameters from these distributions and then simulated a multiple sequence alignment (1000 base pairs) using INDELible

Version 1.03 for each of the gene trees [3]. Note that GTR base frequences (A, C, G, T) were drawn from Dirchlet(113.48869, 69.02545, 78.66144, 99.83793), GTR substitution rates (AC, AG, AT, CG, CT, GT) were drawn from Dirchlet(12.776722, 20.869581, 5.647810 9.863668, 30.679899, 3.199725), and $\alpha$ was drawn from LogNormal(-0.470703916, 0.348667224), where the first parameter is the meanlog and the second parameter is the sdlog.

#### 3.3 Gene Tree Estimation

On gene trees with four or more species, we estimated gene trees using RAxML Version 8.2.12 with the command:

```
raxmlHPC-SSE3 -m GTRGAMMA -p <random seed> -n <output name> -s <alignment file>
```

Sequences were truncated to the first 25, 50, 100, and 250 nucleotides to produce datasets with varying levels of gene tree estimation error (GTEE). GTEE was measured by the normalized RF distance between the true and the estimated gene family trees. Sequence lengths of 25, 50, 100, and 250 resulted in average GTEE of 67%, 52%, 35%, and 19%, respectively.

#### 3.4 Species Tree Estimation Commands

We estimated species trees for each data set using either the first 25, 50, 100, or 500 gene trees; recall that gene trees were either true or had one of four GTEE levels. Gene trees were formatted for each species tree method with custom Python scripts using Dendropy [9].

FastMulRFS operates in three steps. First, the gene family trees are preprocessed to make them singly-labeled trees (Algorithm 1) using a custom Python script. Then, ASTRAL is used to generate the constrained search space $\Sigma$ of allowed bipartitions; we used ASTRAL Version 5.6.3. Finally, the set of singly-labeled trees and constrained search space $\Sigma$ are passed to FastRFS, which returns the optimal trees. The commands for FastRFS and ASTRAL are given below.

ASTRAL-multi and FastMulRFS both use ASTRAL version 5.6.3, with the same command:

```
java -Xms2000M -Xmx20000M -jar astral.5.6.3.jar -i <gene tree file> \
    -a <species to gene name map file> -o <output file> &> <log file>
```

DupTree (download: `http://genome.cs.iastate.edu/CBL/DupTree/linux-i386.tar.gz`) was run with the command:

```
./duptree -i <gene tree file> -o <output file> &> <log file>
```

MulRF Version 2.1 was run with the command:

```
./MulRFSupertreeLin -i <gene tree file> -o <output file> &> <log file>
```

FastRFS Version 1.0 (Initial Release) was run with the command:

```
./FastRFS -i <gene tree file> -o <output file> &> <log file>
```

Table S1: For each dataset, we computed the mean (± standard deviation), the minimum, and the maximum number of copies of each species across all gene tree. We then averaged these values across all species, and then across all ten replicate datasets in order to report a single value per model condition.

| GDL rate | ILS (AD%) | Mean ± Standard deviation | Minimum | Maximum |
|---|---|---|---|---|
| *Species tree height:* $1.8 \times 10^9$ *generations (assumes* 10 *generation per year)* | | | | |
| $1 \times 10^{-10}$ | 2% | 1.0 ±0.6 | 0 | 4.4 |
| $1 \times 10^{-10}$ | 12% | 1.0 ±0.6 | 0 | 4.3 |
| $2 \times 10^{-10}$ | 2% | 1.0 ± 0.9 | 0 | 6.0 |
| $2 \times 10^{-10}$ | 12% | 1.0 ± 0.9 | 0 | 5.9 |
| $5 \times 10^{-10}$ | 2% | 1.0 ± 1.3 | 0 | 9.6 |
| $5 \times 10^{-10}$ | 12% | 1.0 ± 1.4 | 0 | 9.6 |
| *Species tree height:* $2 \times 10^8$ *generations (assumes* $1.\bar{1}$ *generation per year)* | | | | |
| $1 \times 10^{-10}$ | 23% | 1.0 ±0.2 | 0 | 2.3 |
| $1 \times 10^{-10}$ | 22% | 1.0 ±0.2 | 0 | 2.3 |
| $2 \times 10^{-10}$ | 22% | 1.0 ± 0.3 | 0 | 2.8 |
| $2 \times 10^{-10}$ | 64% | 1.0 ± 0.3 | 0 | 2.9 |
| $5 \times 10^{-10}$ | 64% | 1.0 ± 0.5 | 0 | 3.5 |
| $5 \times 10^{-10}$ | 63% | 1.0 ± 0.5 | 0 | 3.6 |

Table S2: For the fungal biological dataset, we computed the mean (± standard deviation), the minimum, and the maximum number of copies of each species across 5,351 gene trees. We also computed the number of gene trees with more than 1, 2, 5, 10, and 20 copies of a species (i.e., > 1 indicates the number of gene trees out of 5,351 with more than 1 copy of the species.)

| Species | Mean ± Std | Minimum | Maximum | >1 | >2 | >5 | >10 | >20 |
|---|---|---|---|---|---|---|---|---|
| A. gossypii | 0.85 ± 0.58 | 0 | 13 | 267 | 41 | 4 | 1 | 0 |
| C. alibicans | 1.04 ± 0.65 | 0 | 7 | 596 | 145 | 6 | 0 | 0 |
| C. glabrata | 0.93 ± 0.81 | 0 | 27 | 589 | 104 | 9 | 4 | 1 |
| C. guilliermondii | 0.99 ± 0.70 | 0 | 11 | 587 | 143 | 7 | 1 | 0 |
| C. lusitaniae | 0.95 ± 0.62 | 0 | 10 | 523 | 101 | 2 | 0 | 0 |
| C. parapsilosis | 1.00 ± 0.73 | 0 | 12 | 588 | 148 | 15 | 2 | 0 |
| C. tropicalis | 1.04 ± 0.73 | 0 | 8 | 655 | 174 | 17 | 0 | 0 |
| D. hansenii | 1.02 ± 0.65 | 0 | 7 | 588 | 138 | 10 | 0 | 0 |
| K. latics | 0.89 ± 0.63 | 0 | 15 | 336 | 55 | 10 | 2 | 0 |
| K. waltii | 0.88 ± 0.71 | 0 | 18 | 315 | 57 | 12 | 4 | 0 |
| L. elongisporus | 0.98 ± 0.66 | 0 | 9 | 576 | 115 | 7 | 0 | 0 |
| S. bayanus | 0.94 ± 0.81 | 0 | 23 | 684 | 117 | 9 | 2 | 1 |
| S. castellii | 1.01 ± 0.91 | 0 | 25 | 826 | 148 | 17 | 6 | 1 |
| S. cerevisiae | 1.03 ± 1.09 | 0 | 42 | 752 | 146 | 19 | 5 | 3 |
| S. mikatae | 0.92 ± 0.76 | 0 | 18 | 629 | 114 | 11 | 2 | 0 |
| S. paradoxus | 0.95 ± 0.83 | 0 | 25 | 690 | 125 | 13 | 2 | 1 |

### 4 Results on Simulated Datasets

#### 4.1 Results with DupTree

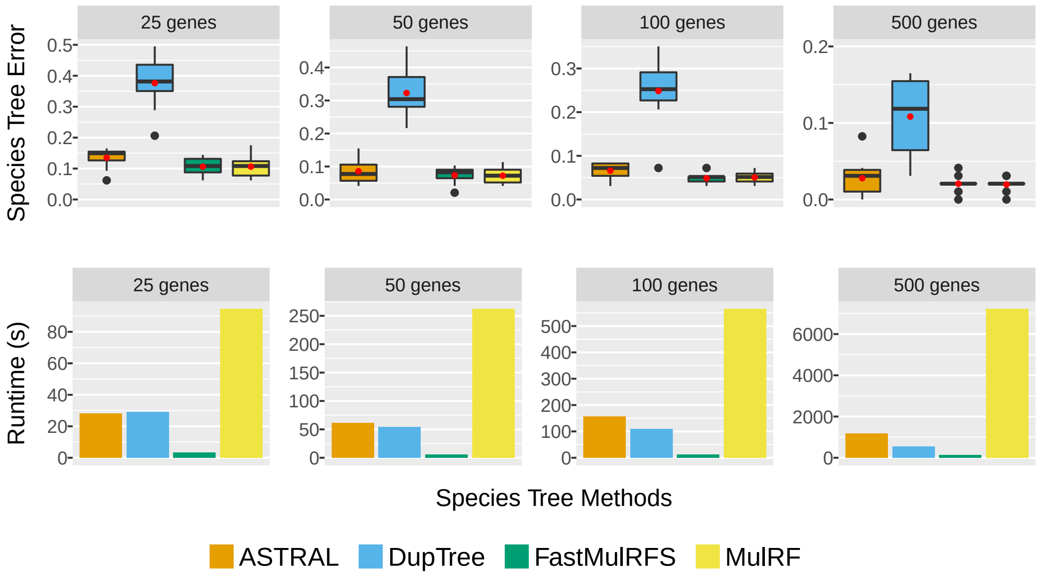

Figure S2: Species tree error and running time (seconds) are shown for FastMulRFS, MulRF, ASTRAL-multi, and DupTree under the easier model conditions, each with 100 species. The model conditions have substantial GTEE (52%), low GDL (D/L rate: $1 \times 10^7$), and very low ILS (2%). Red dots (first row of each subfigure) and bars (second row of each subfigure) are means for 10 replicates.

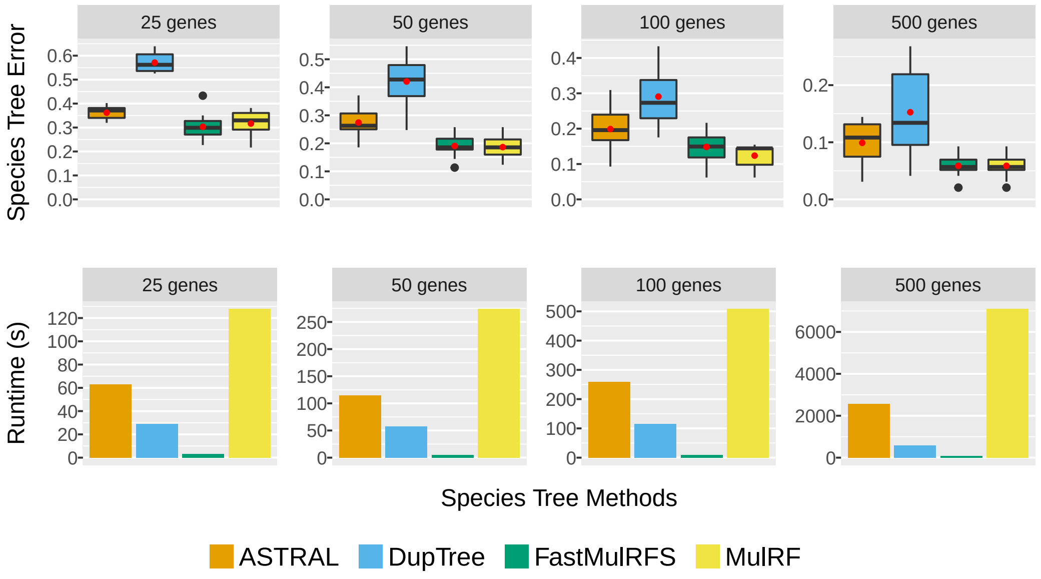

Figure S3: Species tree error and running time (seconds) are shown for FastMulRFS, MulRF, ASTRAL-multi, and DupTree, under the harder model conditions, each with 100 species. The model conditions have substantial GTEE (52%), high GDL (D/L rate: $5 \times 10^7$), and moderate ILS (12%). Red dots (first row of each subfigure) and bars (second row of each subfigure) are means for 10 replicates.

#### 4.2 RFS-multree Criterion Scores

We compared FastMulRFS and MulRF with respect to RFS-multree criterion scores. Out of the 1200 datasets analyzed:

- FastMulRFS was worse than MulRF on 67 datasets
- FastMulRFS was equal to MulRF on 907 datasets
- FastMulRFS was better than MulRF on 226 datasets

This comparison shows that MulRF was actually quite good at finding good scores, but that when the two methods produced trees with different scores, then FastMulRFS dominated MulRF. We also compare methods in terms of species tree accuracy (Table S3) and running time (Table S4).

#### 4.3 Additional Results for FastMulRFS, MulRF, ASTRAL-multi

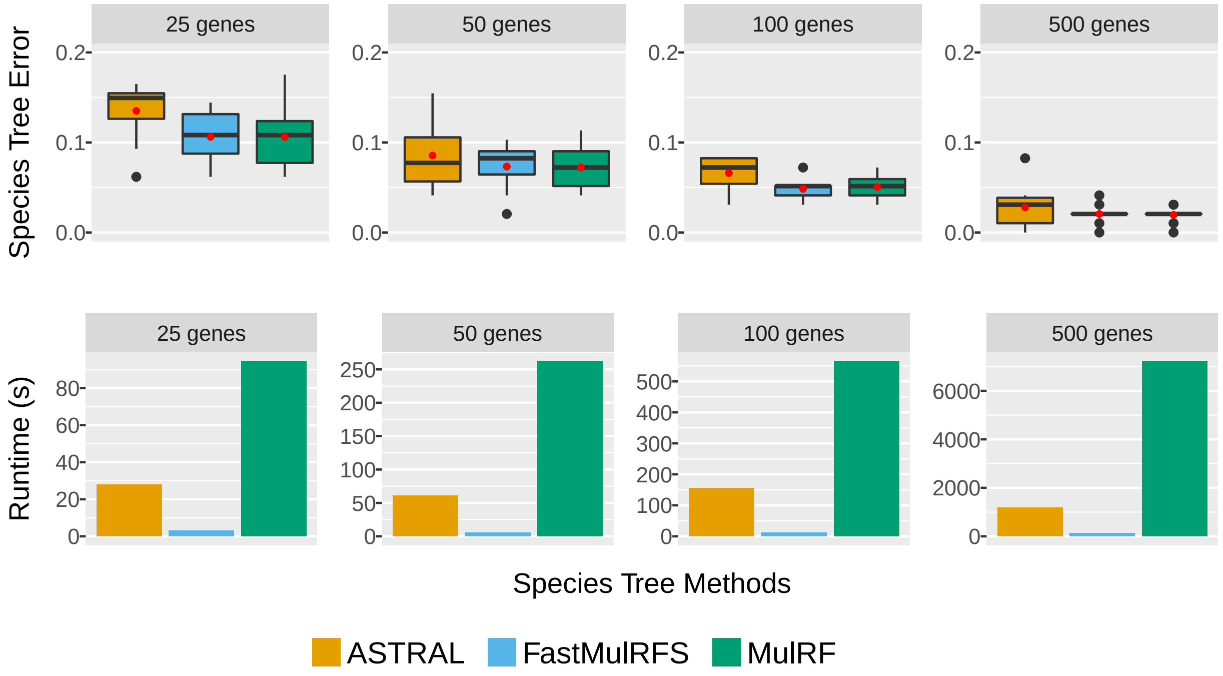

Figure S4: Species tree error (i.e., normalized RF distance) and running time (seconds) are shown for three methods (FastMulRFS, MulRF, and ASTRAL-multi) under the easiest model condition with 100 species. All data sets have substantial GTEE (52%), low GDL (D/L rate: $1 \times 10^7$), and very low ILS (2%). Red dots (first row) and bars (second row) are means for 10 replicate datasets.

Table S3: Species tree error (i.e., normalized RF distance between the true and estimated species trees) averaged across 10 replicate datasets for each of the model conditions on 100-species simulated datasets for ASTRAL-multi / FastMulRFS / MulRF. ILS level is measured using the AD (average normalized RF distance) between true locus trees and true gene trees.

| GTEE | 25 genes | 50 genes | 100 genes | 500 genes |
|---|---|---|---|---|
| *ILS: 2% AD, D/L Rate: 1E-10* | | | | |
| 67% | 0.26 / 0.22 / 0.23 | 0.19 / 0.12 / 0.13 | 0.17 / 0.08 / 0.09 | 0.10 / 0.04 / 0.05 |
| 52% | 0.14 / 0.11 / 0.11 | 0.09 / 0.07 / 0.07 | 0.07 / 0.05 / 0.05 | 0.03 / 0.02 / 0.02 |
| 35% | 0.07 / 0.07 / 0.06 | 0.05 / 0.05 / 0.05 | 0.04 / 0.03 / 0.03 | 0.02 / 0.02 / 0.02 |
| 19% | 0.04 / 0.04 / 0.04 | 0.03 / 0.03 / 0.02 | 0.02 / 0.02 / 0.02 | 0.01 / 0.01 / 0.01 |
| 0% | 0.01 / 0.01 / 0.01 | 0.01 / 0.00 / 0.00 | 0.00 / 0.00 / 0.00 | 0.00 / 0.00 / 0.00 |
| *ILS: 2% AD, D/L Rate: 2E-10* | | | | |
| 67% | 0.33 / 0.30 / 0.36 | 0.27 / 0.21 / 0.21 | 0.19 / 0.14 / 0.16 | 0.12 / 0.06 / 0.07 |
| 52% | 0.19 / 0.16 / 0.16 | 0.15 / 0.10 / 0.10 | 0.11 / 0.08 / 0.08 | 0.05 / 0.04 / 0.04 |
| 35% | 0.11 / 0.10 / 0.09 | 0.08 / 0.05 / 0.05 | 0.05 / 0.04 / 0.04 | 0.03 / 0.02 / 0.02 |
| 19% | 0.07 / 0.05 / 0.05 | 0.04 / 0.03 / 0.03 | 0.03 / 0.03 / 0.03 | 0.01 / 0.01 / 0.01 |
| 0% | 0.01 / 0.01 / 0.01 | 0.01 / 0.00 / 0.00 | 0.01 / 0.00 / 0.00 | 0.01 / 0.00 / 0.00 |
| *ILS: 2% AD, D/L Rate: 5E-10* | | | | |
| 67% | 0.48 / 0.42 / 0.48 | 0.37 / 0.31 / 0.33 | 0.32 / 0.21 / 0.22 | 0.19 / 0.08 / 0.07 |
| 52% | 0.33 / 0.28 / 0.29 | 0.23 / 0.18 / 0.17 | 0.18 / 0.10 / 0.10 | 0.09 / 0.05 / 0.04 |
| 35% | 0.19 / 0.18 / 0.16 | 0.14 / 0.12 / 0.09 | 0.11 / 0.07 / 0.06 | 0.05 / 0.03 / 0.02 |
| 19% | 0.11 / 0.11 / 0.08 | 0.09 / 0.07 / 0.05 | 0.07 / 0.04 / 0.03 | 0.02 / 0.02 / 0.02 |
| 0% | 0.04 / 0.01 / 0.01 | 0.04 / 0.01 / 0.01 | 0.02 / 0.01 / 0.01 | 0.00 / 0.00 / 0.00 |
| *ILS: 12% AD, D/L Rate: 1E-10* | | | | |
| 67% | 0.32 / 0.27 / 0.27 | 0.24 / 0.17 / 0.17 | 0.18 / 0.12 / 0.14 | 0.11 / 0.06 / 0.06 |
| 52% | 0.19 / 0.15 / 0.15 | 0.13 / 0.12 / 0.11 | 0.12 / 0.07 / 0.08 | 0.05 / 0.04 / 0.04 |
| 35% | 0.11 / 0.09 / 0.09 | 0.07 / 0.06 / 0.06 | 0.05 / 0.05 / 0.05 | 0.03 / 0.02 / 0.02 |
| 19% | 0.07 / 0.06 / 0.06 | 0.05 / 0.04 / 0.04 | 0.03 / 0.03 / 0.03 | 0.02 / 0.02 / 0.01 |
| 0% | 0.04 / 0.03 / 0.04 | 0.02 / 0.02 / 0.02 | 0.01 / 0.02 / 0.02 | 0.01 / 0.01 / 0.01 |
| *ILS: 12% AD, D/L Rate: 2E-10* | | | | |
| 67% | 0.35 / 0.30 / 0.34 | 0.26 / 0.22 / 0.23 | 0.20 / 0.14 / 0.15 | 0.11 / 0.08 / 0.07 |
| 52% | 0.19 / 0.16 / 0.16 | 0.15 / 0.12 / 0.12 | 0.12 / 0.08 / 0.07 | 0.06 / 0.03 / 0.03 |
| 35% | 0.12 / 0.10 / 0.10 | 0.09 / 0.09 / 0.08 | 0.08 / 0.06 / 0.05 | 0.03 / 0.02 / 0.02 |
| 19% | 0.07 / 0.06 / 0.06 | 0.05 / 0.05 / 0.05 | 0.05 / 0.04 / 0.03 | 0.02 / 0.02 / 0.02 |
| 0% | 0.04 / 0.03 / 0.03 | 0.03 / 0.03 / 0.02 | 0.02 / 0.01 / 0.01 | 0.01 / 0.01 / 0.01 |
| *ILS: 12% AD, D/L Rate: 5E-10* | | | | |
| 67% | 0.47 / 0.45 / 0.50 | 0.41 / 0.32 / 0.33 | 0.31 / 0.22 / 0.22 | 0.17 / 0.11 / 0.09 |
| 52% | 0.36 / 0.30 / 0.32 | 0.27 / 0.19 / 0.19 | 0.20 / 0.15 / 0.12 | 0.10 / 0.06 / 0.06 |
| 35% | 0.24 / 0.18 / 0.18 | 0.17 / 0.11 / 0.10 | 0.13 / 0.09 / 0.08 | 0.06 / 0.03 / 0.03 |
| 19% | 0.18 / 0.13 / 0.11 | 0.13 / 0.09 / 0.08 | 0.10 / 0.07 / 0.06 | 0.04 / 0.03 / 0.03 |
| 0% | 0.12 / 0.05 / 0.05 | 0.08 / 0.05 / 0.04 | 0.06 / 0.03 / 0.03 | 0.02 / 0.01 / 0.01 |

Table S4: Running time (in seconds) averaged across 10 replicate datasets on 100-species simulated datasets for each of the model conditions for ASTRAL-multi / FastMulRFS / MulRF. ILS level is measured using the AD (average normalized RF distance) between true locus trees and true gene trees.

| GTEE | 25 genes | 50 genes | 100 genes | 500 genes |
|---|---|---|---|---|
| *ILS: 2% AD, D/L Rate: 1E-10* | | | | |
| 67% | 42 / 4 / 113 | 102 / 8 / 263 | 245 / 21 / 614 | 2148 / 296 / 7089 |
| 52% | 28 / 3 / 95 | 61 / 6 / 262 | 156 / 12 / 565 | 1181 / 143 / 7225 |
| 35% | 22 / 3 / 89 | 51 / 9 / 211 | 121 / 8 / 623 | 756 / 72 / 6921 |
| 19% | 19 / 2 / 86 | 37 / 3 / 226 | 86 / 7 / 596 | 641 / 35 / 6727 |
| 0% | 20 / 2 / 99 | 46 / 8 / 228 | 106 / 4 / 582 | 769 / 17 / 6431 |
| *ILS: 2% AD, D/L Rate: 2E-10* | | | | |
| 67% | 52 / 4 / 91 | 123 / 7 / 273 | 306 / 23 / 580 | 2514 / 199 / 6937 |
| 52% | 36 / 5 / 99 | 76 / 5 / 280 | 213 / 10 / 577 | 1357 / 119 / 7225 |
| 35% | 24 / 3 / 92 | 49 / 11 / 254 | 173 / 8 / 636 | 930 / 66 / 6987 |
| 19% | 26 / 2 / 105 | 47 / 10 / 237 | 159 / 6 / 613 | 978 / 44 / 6852 |
| 0% | 26 / 2 / 103 | 55 / 2 / 197 | 157 / 12 / 597 | 829 / 21 / 6827 |
| *ILS: 2% AD, D/L Rate: 5E-10* | | | | |
| 67% | 86 / 4 / 102 | 232 / 7 / 232 | 418 / 14 / 585 | 4495 / 133 / 6884 |
| 52% | 49 / 4 / 114 | 128 / 5 / 691 | 258 / 10 / 692 | 2615 / 83 / 6793 |
| 35% | 40 / 2 / 112 | 79 / 12 / 265 | 195 / 8 / 565 | 1856 / 60 / 6635 |
| 19% | 32 / 8 / 111 | 84 / 3 / 241 | 177 / 6 / 546 | 1288 / 44 / 6645 |
| 0% | 36 / 5 / 110 | 83 / 11 / 215 | 153 / 12 / 542 | 1308 / 25 / 6608 |
| *ILS: 12% AD, D/L Rate: 1E-10* | | | | |
| 67% | 43 / 5 / 112 | 93 / 8 / 239 | 214 / 20 / 680 | 2107 / 274 / 6882 |
| 52% | 29 / 4 / 111 | 63 / 6 / 573 | 133 / 14 / 603 | 1044 / 132 / 7597 |
| 35% | 21 / 3 / 105 | 55 / 5 / 479 | 125 / 14 / 643 | 726 / 69 / 7351 |
| 19% | 22 / 2 / 77 | 49 / 5 / 232 | 111 / 11 / 610 | 673 / 42 / 6976 |
| 0% | 21 / 2 / 83 | 44 / 4 / 318 | 102 / 6 / 572 | 704 / 29 / 6485 |
| *ILS: 12% AD, D/L Rate: 2E-10* | | | | |
| 67% | 57 / 4 / 109 | 130 / 7 / 510 | 302 / 24 / 694 | 3098 / 206 / 7582 |
| 52% | 32 / 10 / 107 | 74 / 14 / 383 | 193 / 12 / 583 | 1767 / 120 / 7085 |
| 35% | 28 / 10 / 103 | 62 / 13 / 223 | 166 / 12 / 672 | 1052 / 61 / 6896 |
| 19% | 22 / 2 / 106 | 53 / 11 / 238 | 182 / 11 / 541 | 836 / 40 / 7189 |
| 0% | 23 / 2 / 104 | 47 / 5 / 279 | 121 / 8 / 586 | 808 / 32 / 6669 |
| *ILS: 12% AD, D/L Rate: 5E-10* | | | | |
| 67% | 105 / 4 / 109 | 205 / 7 / 431 | 469 / 14 / 599 | 4368 / 124 / 6889 |
| 52% | 63 / 3 / 128 | 115 / 5 / 274 | 258 / 10 / 509 | 2572 / 80 / 7107 |
| 35% | 46 / 3 / 112 | 101 / 4 / 204 | 185 / 16 / 647 | 1896 / 69 / 6745 |
| 19% | 37 / 2 / 112 | 82 / 7 / 261 | 193 / 13 / 638 | 1392 / 44 / 6786 |
| 0% | 37 / 7 / 95 | 73 / 3 / 251 | 173 / 5 / 544 | 1402 / 77 / 6746 |
